# Differentiating filter-induced oscillations from physiological stimulation-evoked potentials in intracranial recordings

**DOI:** 10.64898/2026.05.08.723848

**Authors:** Luka Zivkovic, Srdjan Sumarac, David Crompton, William D. Hutchison, Andres M. Lozano, Suneil K. Kalia, Luka Milosevic

## Abstract

**Introduction:** Stimulation-evoked potentials (SEPs), recorded both during and after deep brain stimulation (DBS) surgery, have shown promise for guiding DBS targeting and programming. However, filtering protocols applied to stimulation trains produce an artifact we call a filter-induced oscillation (FIO) which closely mimics physiological SEPs. Hence, we outline the mechanistic origins of this distortion and describe a means of differentiating it from valid SEP activity.

**Methods:** We recorded in 18 patients undergoing DBS surgery targeting the subthalamic nucleus or globus pallidus internus. We stimulated target nuclei with cathode-first (CF) and anode-first (AF) pulses to record native SEPs, and in white matter tracts (null condition). Recordings were subsequently filtered to illustrate FIO. Next, we filtered harmonic frequencies of an artificial stimulation train to demonstrate FIO origins. Finally, FIO was deliberately generated in white matter recordings with a notch filter, and its behaviour contrasted with SEPs during AF and CF stimulation.

**Results:** Filtering stimulation trains produced FIOs that depended on filter order and corner frequency. We also showed that FIO emerges from filter-induced attenuations of harmonic frequencies which compose stimulation trains, producing oscillations of like frequency around pulses. Finally, FIOs reverse in polarity depending on AF or CF stimulation, whereas SEPs do not.

**Conclusions:** Given the potential for widespread adoption of SEPs in DBS targeting and programming, safe analytical protocols are imperative to avoid the induction of processing-related artifacts which can be misinterpreted as biological signals. Here we establish the necessary theory for identifying FIOs and tuning analytical pipelines to avoid their generation.

## Introduction

Deep brain stimulation (DBS) is a neuromodulation therapy used for management of the motor symptoms of Parkinson’s disease (PD),^1^ among other disorders. Primary DBS targets for PD include the subthalamic nucleus (STN)^2^ and globus pallidus internus (GPi).^3^ During DBS surgery, targeting is informed by electrophysiological recordings of local field potentials (LFP) composed of single-neuron spiking as well as synaptic and glial activity.^4^ These signatures can be interpreted and used to guide structure identification and localization through disaggregation of features including spontaneous single-neuron firing rates and the presence or absence of neural oscillations.^5–7^

An additional category of LFP features is stimulation-evoked potentials (SEPs)—transient polarizations in electrical potential following stimulation suggested to be driven by synaptic transmembrane currents.^8–15^ These SEPs are necessarily stimulus-locked and vary in morphology, latency, and frequency.^11^ One notable SEP signature in the STN is “evoked resonant neural activity” (ERNA).^9,10,16–20^ This oscillatory response (thought to be attributable to the recurrent activation of the globus pallidus externus) possesses multiple recurrent peaks occurring at 2-4 ms intervals, exhibits dynamic changes in amplitude, and settles after roughly 10-20 ms.^18,19,21^ Further, the GPi also expresses a non-oscillatory SEP (thought to be attributable to the activation of striatal afferent projections) also exhibiting distinct behaviours;^22,23^ namely, rapid synaptic depression with successive stimuli after a prominent initial peak of a moderate latency (∼4-8 ms). SEPs have also been observed in recordings of cortical activity during subcortical stimulation,^13,24–26^ and have been detected in different structures using stereo-electroencephalography during invasive monitoring for epilepsy.^27,28^ Recent work showing that SEP latencies in the STN align with reductions of instantaneous firing rates corroborates their neural origins.^18^ Furthermore, SEP waveforms match the morphologies and dynamics of postsynaptic currents and potentials in preclinical preparations.^29–31^ As such, SEPs as potential markers of synaptic activations of local network inputs provide an immense opportunity for the development of DBS targeting, postoperative programming, and investigation of therapeutic mechanisms to be informed by the dynamics of underlying network connectivity recorded with impressive resolution and accuracy.^32^

Despite the overall utility of SEPs, analysis of their recordings is complicated by stimulation artifacts^13,15^ in combination with various filtering techniques during post-processing phases of signal acquisition and cleaning. Stimulation artifacts present as large, short-duration changes in amplitude mimicking jump discontinuities in an otherwise relatively stable physiological LFP. In this work, however, we demonstrate that applying filters introduces an artificial oscillation surrounding each impulse which we have termed a “filter-induced oscillation” (FIO). In essence, this behaviour is a filter-derived generalization of the well-known Gibbs’ phenomenon.^33^ The danger of FIOs lies in their ability to resemble SEP signatures which can also manifest as high-frequency oscillatory signals following stimulation impulses. The widespread application of SEPs in the context of DBS targeting, programming, and neurophysiological investigations necessitates safe analytical practices which ensure the integrity of acquired biological signatures and an avoidance of the inadvertent misinterpretation of artifacts. The current literature does not address this commonly-occurring matter directly. Therefore, in this work we introduce FIOs using human intracranial recordings from different subcortical sites with and without known SEP variants, demonstrate the underlying mechanisms which produce FIOs, and apply a stimulation phase-reversal technique to outline a useful means of differentiating physiological SEPs from FIO artifacts.

## Methods

Data were collected from 10 individuals (13 unique sites) undergoing bilateral DBS surgery targeting the STN, and 8 individuals (13 unique sites) undergoing surgery targeting the GPi. Participants provided written informed consent, and all activities were approved by the University Health Network Research Ethics Board.

Two electrically-isolated microelectrodes (spaced 600 μm apart; 100 – 200 kΩ impedance) were advanced through a predefined DBS surgical trajectory, using a common ground and reference. Recordings (10 kHz sampling rate) were acquired using the Cerebus system (Blackrock Neurotech, Salt Lake City, USA). Microstimulation was supplied using a current-controlled, charge-balanced stimulator (CereStim, Blackrock Neurotech).

At 7 of the 13 recording sites within STN DBS surgery, microelectrodes were stationed in the STN whereas the remaining 6 sites were localized to white matter tracts (WMT; between thalamus and STN^5^—no expected SEPs). At the 13 sites recorded from during GPi DBS surgery, microelectrodes were stationed in the GPi only. In all instances, the pair of microelectrodes remained at the same depth. One microelectrode was used for supplying stimulation, while the other recorded the resultant responses. At all STN and WMT sites, we stimulated (monopolar, biphasic, 100 μA; 150 μs pulse-width, 53 μs inter-phase delay) at 100 Hz using a cathode-first (CF) pulse design, followed by an additional train of anode-first (AF) stimulation. For all GPi sites, we stimulated at 1 Hz and 100 Hz (CF and AF for all). 1 Hz stimulation trains were used for GPi sites given the rapidly depressing nature of native SEPs with high stimulation frequencies.^34^

### Demonstrating FIO

To demonstrate FIOs in real recordings, one stimulation train (100 Hz, CF) from each of STN, GPi, and WMT sites (Figure 1A) was processed using a Butterworth highpass filter with varying corner frequencies and filter orders to generate heterogeneous FIOs (Figure 1B). Next, to quantitatively assess overall FIO distortion, we determined the deviation of filtered traces from their raw signals. Raw recordings were first detrended by subtracting the output of a median filter. This nonlinear “moving-median” method is resistant to FIO generation at sufficiently large kernel sizes since sharp-transient deviations in amplitude, like those in stimulation impulses, do not directly contribute to the computation of its final output value at a given time point, but are excluded as nearing the start and end of an ordered sequence used to determine the median of each window. Raw traces were subsequently filtered (Butterworth highpass) using different combinations of filter orders and corner frequencies to induce FIO, and the root-mean-squared error (RMSE) of inter-stimulus intervals (ISI) of each filtered stimulation train was computed against its unfiltered (raw) counterpart. Subsequent RMSE values of each ISI were normalized to the standard deviation of the corresponding raw ISI. Deviation scores for all ISIs in all stimulation trains were then averaged per filter parameter combination.

**Figure 1:**
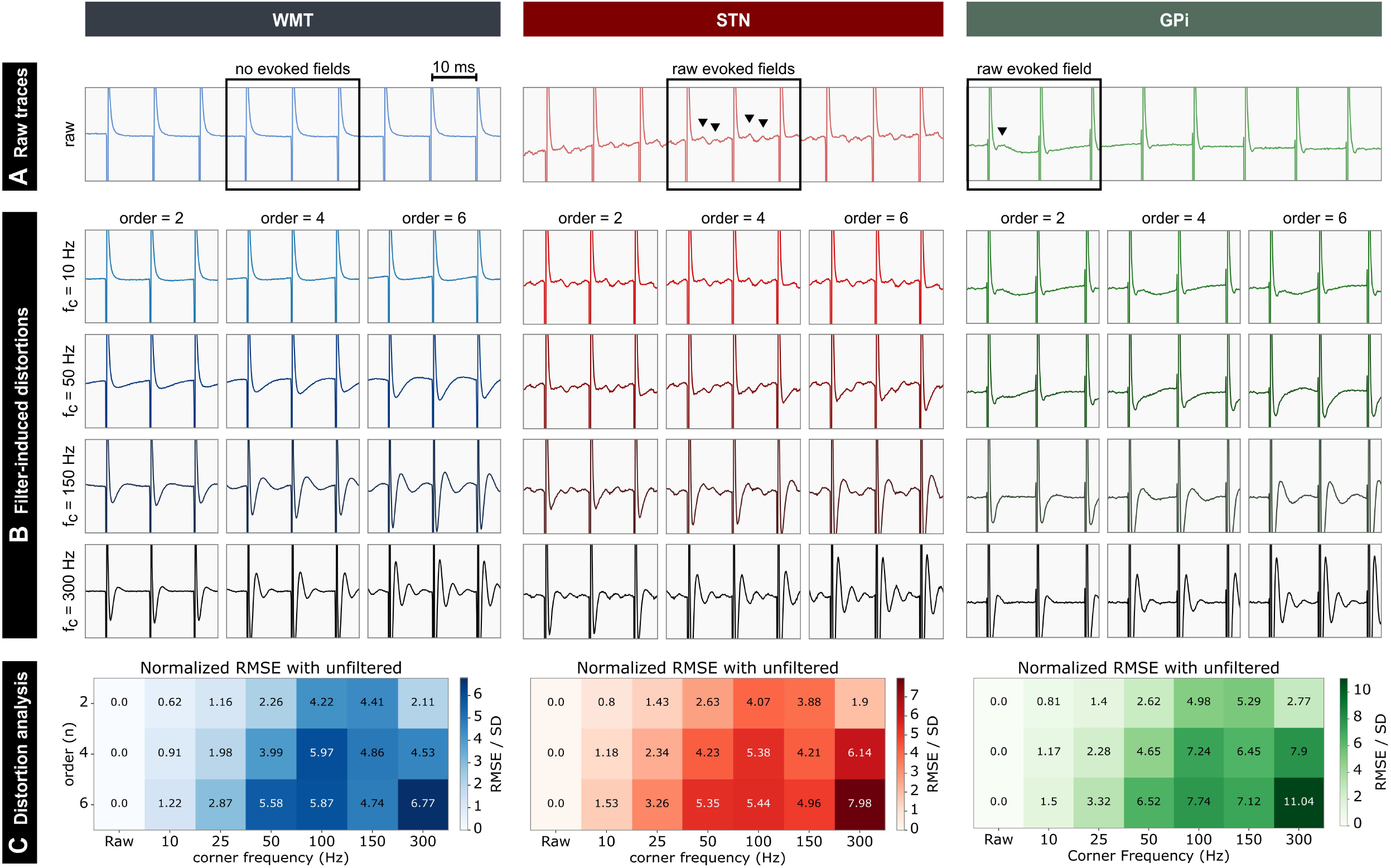
Filter-induced oscillations dependencies, practical dynamics, and filter effects on stimulation trains containing and lacking evoked fields. **A)** Raw traces of stimulation train segments containing exemplary SEP signatures in STN (ERNA) and GPi (non-oscillatory single peak). WMT stimulation contained no evoked fields as a control state. SEP peaks are marked with black triangle arrows for clarity. **B)** The effects of a Butterworth filter at varying orders (*n* = 2, 4, 6) and corner frequencies (*f*_*c*_ = 10, 50, 150, 300 Hz) on the raw stimulation trains, for each category of stimulation/recording sites. Corner frequency is varied along the rows, where orders are varied along columns. **C)** Variance analysis quantifying the deviation of filtered signals from their raw forms using a normalized RMSE at varying filter orders (columns) and corner frequencies (rows). This analysis quantitatively reflects observed distortions in waveform morphologies.

### Mechanisms of FIO generation

We investigated the time and frequency domain effects of various filter types to outline the mechanisms of FIO production in an electrophysiological signal. First, we generated an artificial stimulation train using a simplified equivalent circuit model of recorded stimulation impulses. This consisted of a current source (approximating current-controlled stimulation) and a parallel circuit composed of a capacitor (non-Faradaic effects) and a resistor (Faradaic effects) in series with a battery (*E*_Nernst_ = 0 V), all representing the tissue-electrode interface.^35–37^ Current through the resistor was assumed to be roughly proportional to the sampled voltage in recordings over time. We constructed an input current source to the circuit model with stimulation pulse parameters used in our intracranial recordings (biphasic, cathode-first, 100 μA; 150 μs pulse-width, 53 μs inter-phase delay). Next, we fit our resultant function to a waveform average of the stimulation impulses recorded from a single stimulation train in WMT to obtain a data-informed impulse shape. Next, the resultant stimulation impulse was inserted into a signal of 0 V for all time points for a given duration, and repeated at an interval of 10 ms to generate an artificial 100 Hz stimulation train. This signal was subject to filtering by notch, highpass, lowpass, and bandpass filter designs to demonstrate the generation of FIO. We used this artificial stimulation train for this analysis to simulate an approximation of pure and ideal stimulation artifacts in the absence of any contribution from physiological responses and other sources of noise, in order to systematically characterize FIO.

### Differentiating FIO from stimulation-evoked potentials

To demonstrate a simple means of differentiating FIO artifacts from physiological SEPs, we compared average ISI traces in response to both stimulation phase orders (AF, CF). Since both stimulation phase orders can activate nearby axons and/or their terminals, evoked responses within the signal which are synaptic in nature are expected to retain their peak polarity (representative of hyper-/de-polarizing currents) regardless of stimulation parameters, whereas those sourced in the stimulation artifact are expected to reverse their polarities with the impulse.^13,14,21,38–40^ Within WMT stimulation trains, a 275 Hz FIO was deliberately generated using a notch filter (*f*_c_ = 275 Hz; Q = 9.17) to mimic the approximate frequency of STN ERNA. Next, average waveforms were generated for small segments (10 ms in duration) of all signal variants (WMT, FIO, STN, and GPi) temporally aligned at each stimulation impulse (1 Hz stimulation trains used for GPi sites, 100 Hz stimulation trains used for all others). Mean waveforms were computed and normalized according to the maximum amplitude per stimulation train (excluding stimulation artifacts).

## Results

### Filtering stimulation trains introduces artificial oscillations

ERNA was present in STN recordings, whereas a non-oscillatory SEP was observed in the GPi trace, and no SEPs were found in WMT stimulation trains as expected (Figure 1A). The application of digital filtering yielded distortions whose resultant waveforms were parameter-dependent (Figure 1B). With increasing corner frequency and filter order, deviations from the raw signatures grew, thereby distorting existing SEPs as well as creating artificial oscillations (FIO) of like frequency and amplitude. At very high corner frequencies and orders, the original SEP waveforms (if present) became indiscernible and were replaced with FIOs in STN, GPi, and WMT. Quantitatively, RMSE values, a proxy metric for degree of FIO distortion, confirm these relationships between filter settings and signal distortions (Figure 1C).

In the case of WMT sites which lack SEP signatures, FIOs are observed as the generation of oscillatory peaks where they otherwise did not exist. In STN SEPs, FIO may also present as accentuations of amplitudes of existing ERNA peaks and troughs. At other parameter combinations, the oscillatory phase of the SEP may be shifted closer to the stimulation artifact, presenting the illusion that the peaks have shorter latencies than what appears in raw signatures. With respect to GPi recordings, FIO may also manifest as emphasized amplitudes, latency shifts, but also the apparent attenuation of an existing peak and the generation of new ones.

### Filter-induced oscillations arise as attenuated harmonic frequencies

To demonstrate the mechanistic origins of FIO, we first present the time and frequency domain representations of an artificial pure stimulation train (Figure 2A). The frequency composition includes a vast harmonic distribution with frequency peaks dispersed at an interval equal to that of the stimulation frequency, as well as inter-harmonic peaks between them. Next, following the application of notch filters centered about specific harmonic frequencies, we observe oscillatory distortions of like frequencies in the time domain around stimulation impulses (Figure 2B). The filter-induced attenuation of harmonic frequency components thus appears in the time domain as an oscillation surrounding pulses. Finally, we subjected the artificial stimulation train to highpass, lowpass, and bandpass filtering to demonstrate the emergence of FIO using commonly applied filter designs (Figure 2C).

**Figure 2:**
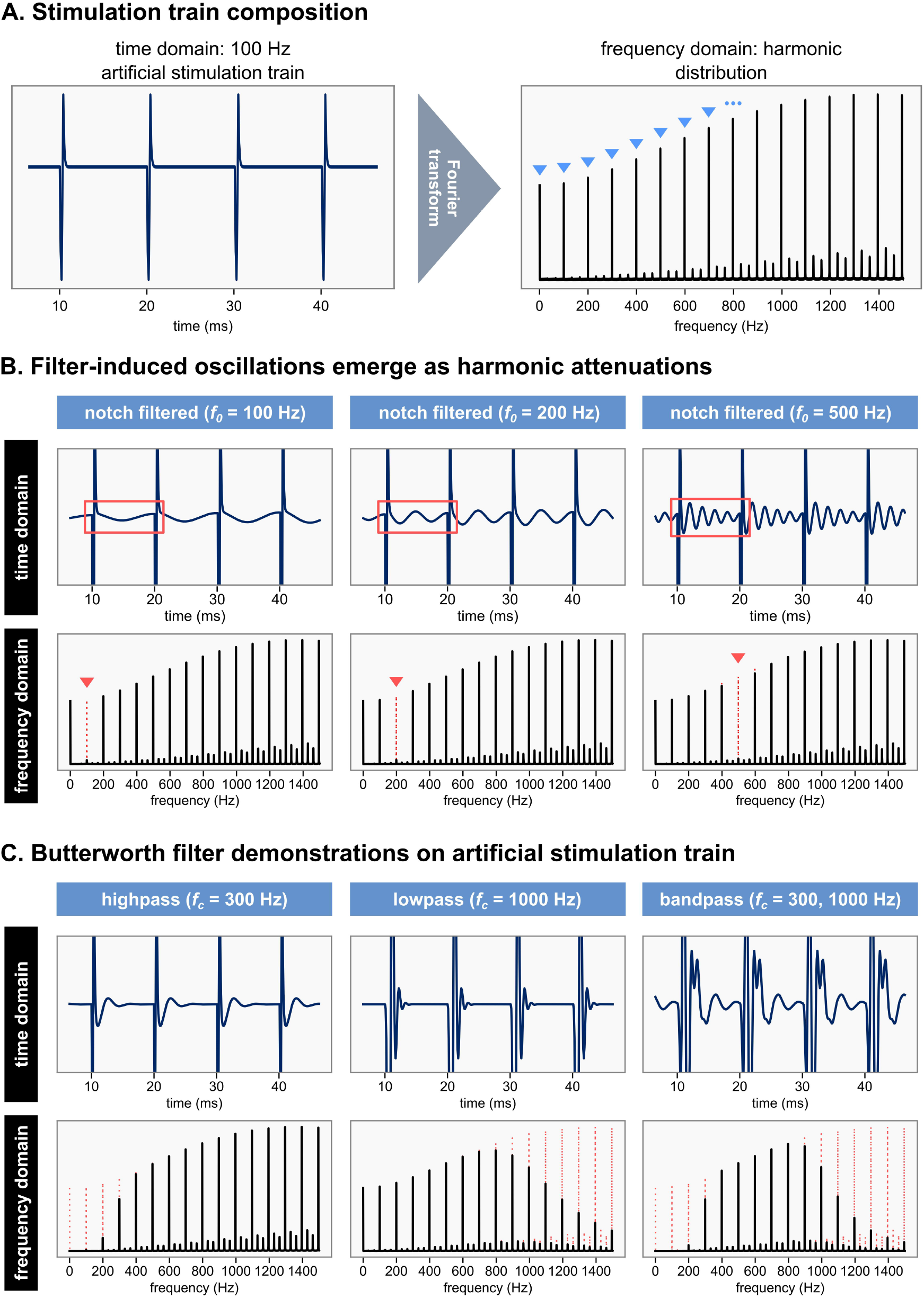
Demonstration of the mechanistic origins of FIO. **A)** An artificial 100 Hz stimulation train and its corresponding frequency domain representation (Fourier transformed signal). The frequency domain shows large harmonic peaks (marked with blue triangle arrows) dispersed at a frequency of 100 Hz (stimulation frequency). **B)** Time and frequency domain outputs of the same artificial stimulation train subjected to notch filtering at 3 distinct frequencies (100, 200, and 500 Hz). Each column contains a distinct center frequency. In the time domain, red boxes outline oscillations equal in frequency to the corresponding notch filter center. In the frequency domain, red triangle arrows point to attenuated harmonic peaks as a result of filtering. **C)** Further demonstrations of the outputs of common Butterworth filter designs on the same artificial stimulation train in time and frequency domains. The time domain shows unique oscillations following impulses, and the frequency representations again show distinct distributions of attenuated harmonic frequencies.

### Differentiating filter-induced oscillations from physiological evoked potentials

Given the capacity for filter techniques to introduce artificial and stimulus-locked deviations in the field potential, we provide a simple means of distinguishing the FIO artifact from valid physiological SEP signatures (Figure 3).

**Figure 3:**
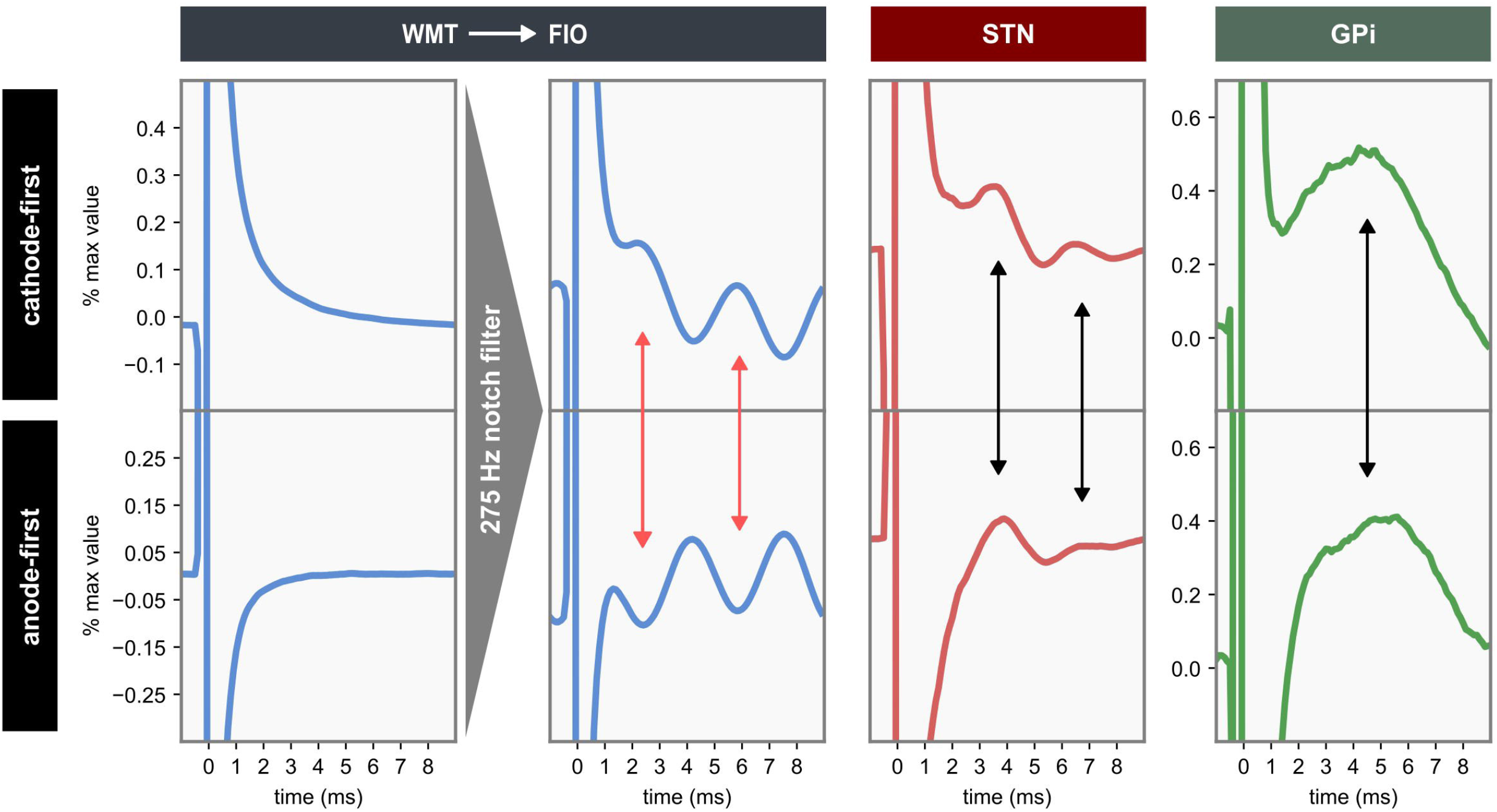
Differentiating a SEP from FIO. A schematic displaying how FIO can be identified through its stimulation-dependent dynamics. Each column and row contain waveform averages of segments centered around all stimulation impulses within trains (1 Hz stimulation for GPi, 100 Hz for remaining sites). WMT recordings (first column) lack SEPs with both AF and CF stimulation. When FIO is introduced (second column) by applying a notch filter with a 275 Hz center frequency, the polarity of the oscillation following the impulse reverses with stimulation phase order. SEP traces in both STN (third column) and GPi (fourth column) do not exhibit this reversal. Instead, STN recordings retain their characteristic fields (ERNA), and so do those in the GPi (non-oscillatory peak). The polarity reversal of FIO is marked by red reciprocal arrows, where peaks in one phase order correspond to troughs in the other. Black reciprocal arrows, however, mark where the polarities remain the same.

Raw WMT waveforms yielded a uniform trace with both CF and AF stimulation, consistent with the lack of any source of SEP or FIO. Where FIO was introduced, a 275 Hz oscillation can be observed with both stimulation configurations. However, the waveforms exhibit opposing polarities, where the oscillation reverses with the stimulation phase order. STN and GPi average waveforms do not exhibit this polarity reversal, but express a constant-phase ERNA and non-oscillatory SEP, respectively. FIO thus exhibits a dependence on the stimulation phase order, whereas unfiltered traces containing valid SEP signatures do not.

## Discussion

In this study, we present observations of FIO signatures in examples of common SEP and null electrophysiological recordings, demonstrate the mechanisms of FIO generation through the time and frequency domain effects of filtering, as well as provide a means of differentiating FIO from valid neural SEPs. These developments establish a cautionary tone in the utilization of SEP signatures in clinical and research practice, emphasizing the use of safe analytical protocols to prevent the study of artifacts as they may mimic valid physiological signals. Based on these findings, the improper application of filtering to stimulation trains may significantly mislead clinicians in their use of SEPs, and result in the reporting of false positive signatures.

The second component of our analysis elucidated the time and frequency domain effects of filtering over stimulation trains. This is a useful means of understanding the mechanistic basis of FIO distortions around impulses. In the frequency domain, the effects of filtering are seen as attenuations of prominent harmonic frequency components which dominate the signal’s construction given their relatively large amplitude. In the time domain, this is the equivalent to the subtraction of sinusoids of equal frequency, amplitude, and phase. Although a filtering operation does not precisely behave in this manner and is instead implemented as some discrete difference equation or a multiplication by a transfer function in the frequency domain, for a practical understanding of FIO generation a filter’s effects can be conceived of as a virtual subtraction of large frequency components constructive of the stimulation artifacts from the relatively stable signal surrounding impulses. Furthermore, the waveform morphologies of FIOs are dependent on filter designs which vary in band type, order, and corner frequencies. We also identified differential behaviour between FIO signatures and neural SEPs, where the former reverses in polarity with an inversion of stimulation phase order in a biphasic pulse, while the latter does not due to its neural origins. Linear filter systems thus, as expected, produced reverse-polarity oscillations around stimulation impulses.

The FIO phenomenon is an instance of a much broader class of “ringing” artifacts which are induced through a variety of means. Similar to our analysis, a study by Hudson and Jones investigated a sister-concept in the context of gamma oscillations in electroencephalography recordings. They found that applying conventional filtering techniques to sharp, high-amplitude, transient noise produces a ringing which can be confused for neural oscillations in the gamma frequency band.^41^ One possible avenue for eliminating the effects of similar sharp-transients like stimulation artifacts is to simply remove them before the application of filtering. This may take the form of algorithmic interpolation or even hardware-based solutions which shunt current away from the amplifier or re-reference to subtract stimulation impulses.^15,42,43^ However, even in these cases, caution is advised as processed traces may contain remnant distortions which repeat regularly at the frequency of stimulation and thus, may produce similar distortions. Alternatively, in cases where highpass filtering is required, one may leave the stimulation artifacts unaltered and instead use detrending methods to remove low frequency components. For example, the low frequency drift may be captured by the output of a median filter (as was used in our analysis) which is highly resistant to the influence of transient, extreme deviations in amplitude as are present in stimulation artifacts. The output of this filter may then itself be smoothed and subsequently subtracted from the raw recording, yielding an effectively highpass filtered signal with no FIO for reasons described in our methods. Otherwise, much caution is advised when applying conventional filtering techniques like a Butterworth approximation, where the choice of safe filter parameters will depend largely on the harmonic distribution of the stimulation train and the desired outcome of the user.

SEPs and other forms of evoked potentials have been increasingly implicated in numerous clinical practices including DBS targeting and programming.^27,44,45^ Their utility as markers for identifying and disaggregating regions of differential afferent types in real time during intracranial and extraoperative recordings further solidifies their potential for widespread adoption.^11,18,46^ Hence, the correct use of processing methods in both clinical, academic, and industry applications is imperative. A failure to account for such nuances may lead to misinterpretations of recorded signatures, improper DBS programming, and inaccurate targeting potentially leading to suboptimal clinical outcomes and later revision.

## Conclusions

In summary, we demonstrated our observations and mechanistic origins of the FIO phenomenon which has the capacity to resemble SEP signatures during neural stimulation and recording. FIO can be distinguished from valid neural activity via differential behaviours revealed through the reversal of stimulation phase order. Moreover, our work reinforces a necessary cautionary sentiment to be had within signal processing phases of analyses and prompts further investigations into safe filtering methods which address many of the considerations we have outlined.

## Data availability

The recorded data and analysis pipeline have been made available in a public repository (https://doi.org/10.5281/zenodo.19140963).

## Acknowledgements

This project has been made possible with the financial support from the Natural Sciences and Engineering Research Council (NSERC) RGPIN-2022-05181 (L.M.); Canadian Institute for Health Research (CIHR) PJT 191880 (L.M., S.K.K.); Mitacs Accelerate Program (L.Z.); Blackrock Neurotech (L.Z.). The authors would like to thank the participants for their invaluable contributions to the work.

